# Characterization of the immune resistance of SARS-CoV-2 Mu variant and the immunity induced by Mu infection

**DOI:** 10.1101/2021.11.23.469770

**Authors:** Keiya Uriu, Paúl Cárdenas, Erika Muñoz, Veronica Barragan, Yusuke Kosugi, Kotaro Shirakawa, Akifumi Takaori-Kondo, Ecuador-COVID19 Consortium, The Genotype to Phenotype Japan (G2P-Japan) Consortium, Kei Sato

## Abstract

We have revealed that the SARS-CoV-2 Mu variant is highly resistant to COVID-19 convalescent sera and vaccine sera.^1^ However, it remains unclear how the immune resistance of the Mu variant is determined. Also, although the Mu variant is highly resistant to the sera obtained from COVID-19 convalescent during early pandemic (i.e., infected with prototypic virus) and vaccinated individuals (i.e., immunized based on prototypic virus), it was unaddressed how the convalescent sera from Mu-infected individuals function. In this study, we revealed that the two mutations in the spike protein of Mu variant, YY144-145TSN and E484K, are responsible for the potent immune resistance of Mu variant. Additionally, we showed that the convalescent sera obtained from the Mu-infected individuals can be broadly antiviral against the Mu variant as well as other SARS-CoV-2 variants of concern/interest. Our findings suggest that developing novel vaccines based on the Mu variant can be more effective against a broad range of SARS-CoV-2 variants.

## Text

As of October 2021, the WHO defined four variants of concern and two variants of interest.^2^ Mu variant represents the most recently recognized variant of interest and has spread mainly in some South American countries such as Colombia and Ecuador.^2^ We have recently revealed that the Mu variant shows a pronounced resistance to antibodies elicited by natural severe acute respiratory syndrome coronavirus 2 (SARS-CoV-2) infection and the BNT162b2 vaccine.^1^ The majority of Mu variants harbor the T95I and YY144-145TSN mutations in the N-terminal domain; the R346K, E484K, and N501Y mutations in the receptor-binding domain; and the D614G, P681H, and D950N mutations in other regions of the spike protein (**Fig. 1A**). However, it remains unclear which mutations determine the pronounced resistance of Mu variant to antiviral sera. To address this, we generated a series of pseudoviruses that harbors the spike protein of the D614G-bearing B.1 lineage virus (parental virus) bearing with each mutation in the Mu variant. Virus neutralization assay was performed with the use of serum samples obtained from 15 coronavirus disease 19 convalescents who were infected early in the pandemic (April 2020) (**Table S1**) and 14 persons who had received the BNT162b2 vaccine (**Table S2**). As shown in **Fig. 1B** (top), two mutations, YY144-145TSN and E484K, conferred the resistance to antibodies induced by natural SARS-CoV-2 infection and vaccination. To verify the effect of these two mutations on the neutralization resistance, we next generated a series of Mu-based pseudoviruses that lose respective mutations. Consistent with the gain-of-function experiments based on the parental virus (**Fig. 1B**, top), the loss-of-function experiments showed that the spike proteins of Mu variant reverting YY144-145TSN or E484K mutations loses the neutralization resistance (**Fig. 1B**, bottom). Since the Mu pseudovirus derivative that loses both YY144-145TSN and E484K mutations almost completely lost the neutralization resistance (**Fig. 1B**, bottom), our data suggest that the pronounced resistance of Mu variant against neutralizing antibodies is attributed to these two mutations.

**Figure 1.**
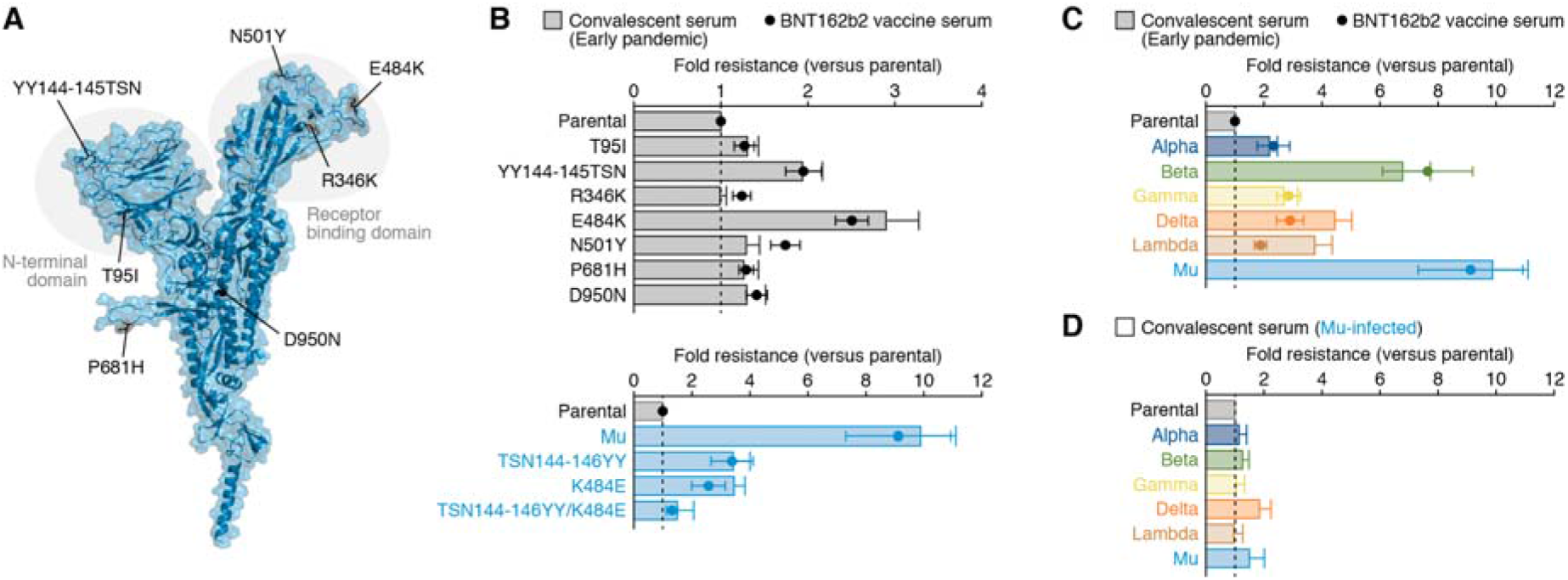
Characterization of the immune resistance of the Mu variant. Panel A shows the position of the mutations in Mu variant. Cartoon and surface models are overlayed. The mutations in Mu variant are indicated. The structure of N-terminal domain is shown in **Fig. S1** in the Supplementary Appendix. Panels B to D show the results of virus neutralization assays. Neutralization assays were performed with the use of pseudoviruses harboring the SARS-CoV-2 spike proteins of parental virus (the B.1 lineage virus, which harbors the D614G mutation)-based derivatives (Panel B, top), the spike proteins of Mu-based derivatives (Panel B, bottom), or the spike proteins of the Alpha, Beta, Gamma, Delta, Lambda or Mu variants (Panels C and D). Serum samples were obtained from 15 convalescent persons who had infected with SARS-CoV-2 in the early pandemic, 14 persons who had received the BNT162b2 vaccine, and 4 convalescent persons who had infected with SARS-CoV-2 Mu variant. In Panels B and C, the heights of the bars (serum samples obtained from the convalescent persons who had infected with SARS-CoV-2 in the early pandemic) and the circles (serum samples obtained from the persons who had received the BNT162b2 vaccine) indicate the average difference in neutralization resistance of the indicated variants as compared with that of the parental virus. In Panel D, the heights of the bars (serum samples obtained from the convalescent persons who had infected with Mu variant) indicate the average difference in neutralization resistance of the indicated variants as compared with that of the parental virus. The error bars indicate standard error of the mean. The vertical dashed lines indicate value 1. The assay of each serum sample was performed in triplicate to determine the 50% neutralization titer. The raw data of the 50% neutralization titer are summarized in **Fig. S2** and **Tables S1-S3** in the Supplementary Appendix. The information regarding the convalescent donors (sex, age, and dates of testing and sampling) and vaccinated donors (sex, age, and dates of second vaccination and sampling) of serum samples are summarized in **Tables S1-S3** in the Supplementary Appendix.

We next assessed the immunological spectrum of the serum samples obtained from the convalescents who had infected with Mu variant (**Table S3**). Although the Mu variant was more than 9 times resistant to the sera induced by natural SARS-CoV-2 infection during early pandemic and vaccination, which is consistent with our recent report (**Fig. 1C**),^1^ the Mu variant did not exhibit resistance to the sera induced by Mu infection (**Fig. 1D**). Notably, the sera induced by Mu infection exhibited broad antiviral effect against various variants of concern/interest (**Fig. 1D**). Altogether, our findings suggest that the use of Mu sequence can be a good strategy for the next-generation vaccine development.

