## Supplementary appendix for "Characterization of the immune resistance of SARS-CoV-2 Mu variant and the immunity induced by Mu infection"

**Table of Contents**

**Contents**	Page

**Materials and Methods**	2-4

Ethics statement

Human sera

Protein homology model

Viral genome sequences

Cell culture

Plasmid construction

Neutralization assay

**Figure S1.** Structure of the N-terminal domain of the spike protein of the Mu variant.	5

**Figure S2.** Summary of the 50% neutralization titer of each serum.	6-7

**Table S1.** Summary of the serum samples obtained from COVID-19 convalescents

who had infected in the early pandemic used in this study.	8-11

**Table S2.** Summary of the serum samples obtained from the persons who had

received the BNT162b2 vaccine used in this study.	12-15

**Table S3.** Summary of the serum samples obtained from COVID-19 convalescents

who had infected with Mu variant used in this study.	16-17

**Table S4.** Primers used for the construction of SARS-CoV-2 spike protein derivatives.	18

**Consortia**	19-20

**Acknowledgments**	21

**Supplemental References**	22

**Materials and Methods**

**Ethics statement**

For the use of human specimens, all protocols involving human subjects recruited at Kyoto University and Universidad San Francisco de Quito were reviewed and approved by the Institutional Review Boards of The Institute of Medical Science, The University of Tokyo (approval ID: 2021-1-0416), Kyoto University (approval ID: G0697), Universidad San Francisco de Quito (approval ID: CEISH P2020-022IN), and the Ecuadorian Ministry of Health (approval IDs: MSP-CGDES-2020-0121-O and MSP-CGDES-061-2020). The export of sera from Ecuador to Japan was approved by ARCSA ID: ARCSA-ARCSA-CGTC-DTRSNSOYA-2021-1626-M. All human subjects provided written informed consent.

**Human sera**

Fifteen serum samples obtained from COVID-19 convalescents who had infected with SARS-CoV-2 in the early pandemic were purchased from RayBiotech (**Table S1**). Peripheral blood was collected four weeks after the second vaccination with BNT162b2 (Pfizer-BioNTech), and the sera of fourteen vaccinated-individuals were isolated (**Table S2**). Peripheral blood was collected from four COVID-19 convalescents who had infected with SARS-CoV-2 Mu variant and the sera were isolated (**Tables S3**). Sera were inactivated at 56°C for 30 min and stored at –80°C until use.

**Protein homology model**

All protein structural analyses were performed using Discovery Studio 2021 (Dassault Systèmes BIOVIA). In **Fig. 1A** and **Fig. S1**, the crystal structure of SARS-CoV-2 spike protein (B.1 lineage; PDB: 7KRS)^3^ was used as the template, and 40 homology models of the spike protein of Mu variant were generated using Build Homology Model protocol MODELLER v9.24.^4^ Evaluation of the homology models was performed using PDF total scores and DOPE scores and the best model for the spike protein of Mu variant was selected.

**Viral genome sequences**

RNA was extracted from the nasopharyngeal swabs of COVID-19 patients using SV total RNA isolation system (Cat# Z3101, Promega). Reverse transcription was performed following the ARTIC protocol SARS-CoV-2 primer scheme v3 (<https://www.protocols.io/view/ncov-2019-sequencing-protocol-bbmuik6w>). The reverse transcription products were purified using AMPure XP magnetic beads (Cat# A63880, Beckman Coulter) according to the manufacturer instructions and then quantified using Qubit RNA assay kit (Cat# Q32852, Thermo Fisher Scientific). Genomic library was generated using the native barcoding expansion 96 kit (Cat# EXP-NBD196, Oxford Nanopore Technologies) with ligation sequencing kit (Cat# LSK-109, Oxford Nanopore Technologies) and loaded into the MinION flow cell (Cat# FLO-MIN 106, Oxford Nanopore Technologies). RAMPART software v1.0.5 from the ARTIC Network (<https://github.com/artic-network/rampart>) was used to monitor the sequencing in real-time. Porechop v0.2.4 (<https://github.com/rrwick/Porechop>) was used to carry out demultiplexing and adapter removal. The ARTIC Network bioinformatics pipeline was employed to create consensus sequences and variant calls (<https://www.protocols.io/view/ncov-2019-sequencing-protocol-bbmuik6w>). To generate the consensus genomes, the reads were mapped on the reference genome of SARS-CoV-2 strain Wuhan-Hu-1 (GenBank accession number MN908947). The sequences were uploaded to the online tool to determine genome clades and identify mutations. Pangolin COVID-19 lineage assigner v.3.1.11^5^ and NextClade v.1.6.0^6^ were used for the lineage classification**.** The viral sequences were deposited in the GISAID database (<https://www.gisaid.org>), and the GISAID IDs are listed in **Table S3**.

**Cell culture**

HEK293T cells (a human embryonic kidney cell line; ATCC CRL-3216) and HOS-ACE2/TMPRSS2 cells,^7^^,^^8^ a derivative of HOS cells (a human osteosarcoma cell line; ATCC CRL-1543) stably expressing human ACE2 and TMPRSS2, were maintained in Dulbecco’s modified Eagle's medium (high glucose) (Wako, Cat# 044-29765) containing 10% fetal calf serum, 100 units penicillin and 100 ug/ml streptomycin.

**Plasmid construction**

Plasmids expressing the SARS-CoV-2 spike proteins of the parental D614G (B.1), Alpha (B.1.1.7), Beta (B.1.351), Gamma (P.1), Delta (B.1.617.2), Lambda (C.37) and Mu (B.1.621) variants were prepared in our previous studies.^8-11^ Plasmids expressing the SARS-CoV-2 spike protein derivatives of parental D614G (B.1) and Mu (B.1.621) variant were generated by site-directed overlap extension PCR using the expression plasmid of either parental D614G (B.1)^8^ and Mu (B.1.621) variant^11^ as the template and the primers listed in **Table S4**. The resulting PCR fragment was digested with KpnI and NotI and inserted into the corresponding site of the pCAGGS vector.^12^ Nucleotide sequences were determined by DNA sequencing services (Eurofins), and the sequence data were analyzed by Sequencher v5.1 software (Gene Codes Corporation).

**Neutralization assay**

Pseudoviruses were prepared as previously described.^8-11^^,^^13^ Briefly, lentivirus (HIV-1)-based, luciferase-expressing reporter viruses were pseudotyped with the SARS-CoV-2 spikes. HEK293T cells (1 × 10^6^ cells) were cotransfected with 1 μg psPAX2-IN/HiBiT,^14^ 1 μg pWPI-Luc2,^14^ and 500 ng plasmids expressing parental S or its derivatives using PEI Max (Polysciences, Cat# 24765-1) according to the manufacturer's protocol. Two days post transfection, the culture supernatants were harvested and centrifuged. The pseudoviruses were stored at –80°C until use.

Neutralization assays were performed as previously described.^9^^,^^11^^,^^13^ Briefly, the SARS-CoV-2 spike pseudoviruses (counting ~20,000 relative light units) were incubated with serially diluted (40-, 120-, 360-, 1,080-, 3,240-, 9,720-, and 29,160-fold dilution at the final concentration) heat-inactivated human sera at 37°C for 1 h. Pseudoviruses without sera were included as controls. Then, an 80 μl mixture of pseudovirus and serum was added to HOS-ACE2/TMPRSS2 cells (10,000 cells/50 μl) in a 96-well white plate. Two days post infection, the infected cells were lysed with a One-Glo luciferase assay system (Promega, Cat# E6130), and the luminescent signal was measured using a GloMax explorer multimode microplate reader 3500 (Promega). The assay of each serum was performed in triplicate, and the 50% neutralization titer was calculated using Prism 9 (GraphPad Software).

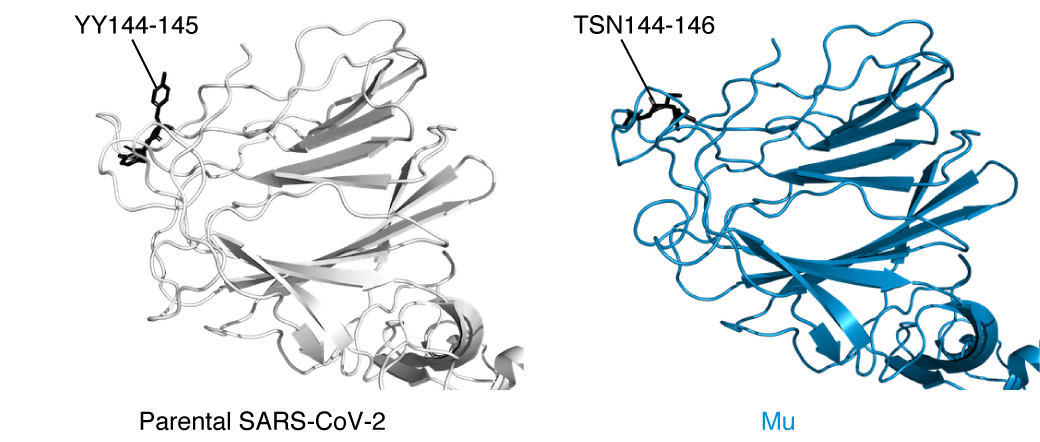

**Figure S1. Structure of the N-terminal domain of the spike protein of the Mu variant.** The N-terminal domain of the crystal structure of SARS-CoV-2 spike protein (PDB: 7KRS, left) and a homology model of Mu variant (right) are shown. The YY144-145TSN mutation in the Mu variant (TSN144-146) and relevant residues in the parental spike protein (YY144-145) are indicated in the panel.

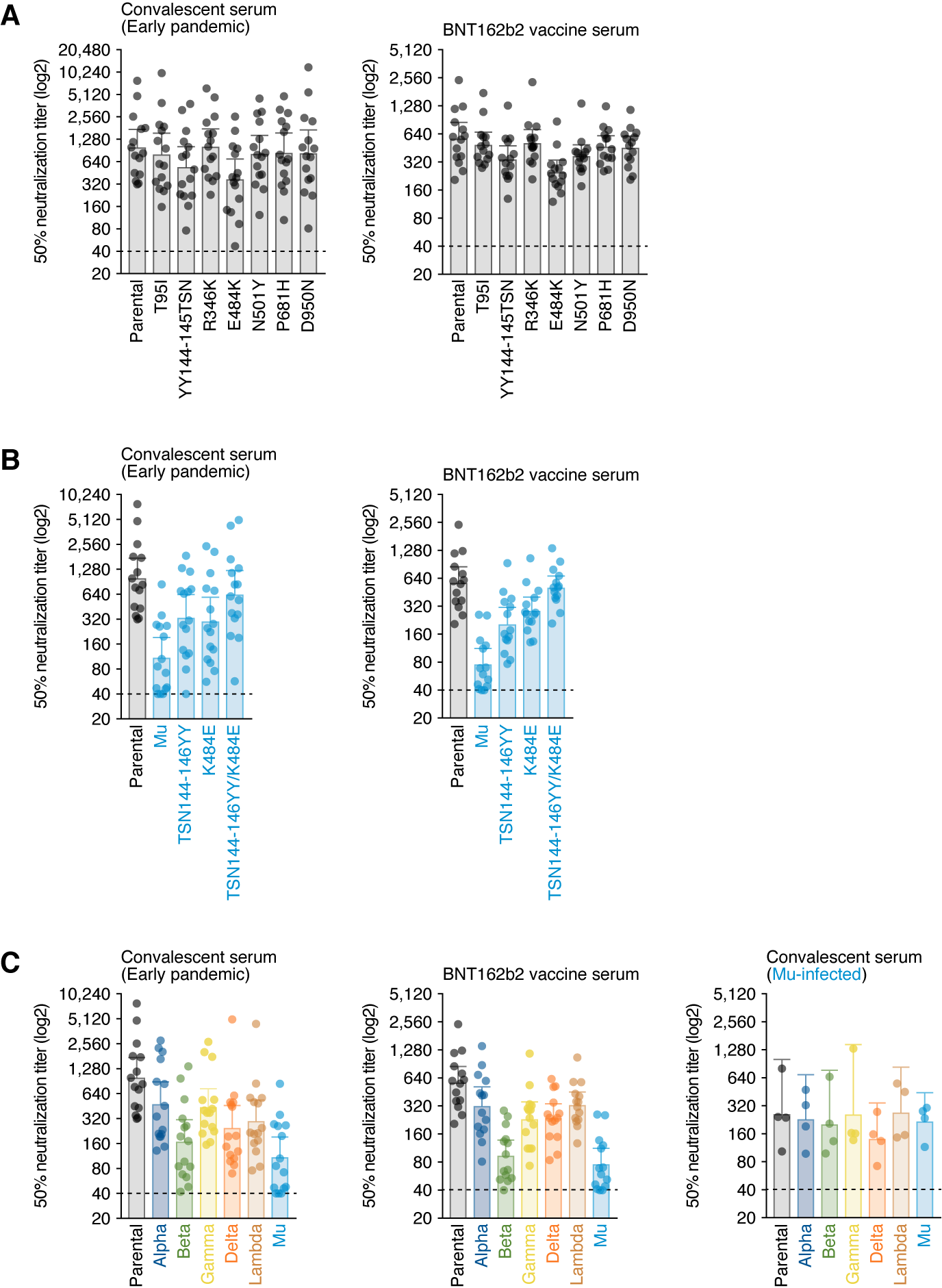

**Figure S2. Summary of the 50% neutralization titer of each serum.** Neutralization assays were performed with the use of pseudoviruses harboring the SARS-CoV-2 spike proteins of parental virus (the B.1 lineage virus, which harbors the D614G mutation)-based derivatives (**A**), the spike proteins of Mu-based derivatives (**B**), or the spike proteins of the Alpha, Beta, Gamma, Delta, Lambda or Mu variants (**C**). Serum samples were obtained from 15 convalescent persons who had infected with SARS-CoV-2 in the early pandemic (**A–C**), 14 persons who had received the BNT162b2 vaccine (**A–C**), and 4 convalescent persons who had infected with SARS-CoV-2 Mu variant (**C**). The assay of each serum sample was performed in triplicate to determine the 50% neutralization titer. Each data point represents an individual sample (circles) and indicates the 50% neutralization titer obtained with each sample against the indicated pseudovirus. The heights of the bars and the numbers over the bars indicate the geometric mean titers, and the I bars indicate 95% confidence intervals. The horizontal dashed lines indicate the limit of detection. The average difference in neutralization resistance of the indicated variants as compared with that of the parental virus is summarized in **Fig. 1**. The assay of each serum sample was performed in triplicate to determine the 50% neutralization titer. The raw data of the 50% neutralization titer and the information regarding the convalescent donors (sex, age, and dates of testing and sampling) and vaccinated donors (sex, age, and dates of second vaccination and sampling) of serum samples are summarized in **Tables S1-S3**.

**Consortia**

**The Genotype to Phenotype Japan (G2P-Japan) Consortium**

**The Institute of Medical Science, The University of Tokyo, Japan**

Jumpei Ito, Daichi Yamasoba, Izumi Kimura, Mai Suganami, Akiko Oide,

Miyabishara Yokoyama, Mika Chiba

**Tokai University, Japan**

So Nakagawa, Jiaqi Wu, Miyoko Takahashi

**Kyoto University, Japan**

Yasuhiro Kazuma, Ryosuke Nomura, Yoshihito Horisawa, Kayoko Nagata, Yohei Yanagida,

Yugo Kawai, Yusuke Tashiro

**Chiba University, Japan**

Atsushi Kaneda, Taka-aki Nakada, Motoaki Seki, Ryoji Fujiki, Tadanaga Shimada,

Kiyoshi Hirahara, Koutaro Yokote, Toshinori Nakayama

**Hiroshima University, Japan**

Takashi Irie, Ryoko Kawabata, Nanami Morizako

**Hokkaido University, Japan**

Takasuke Fukuhara, Kenta Shimizu, Kana Tsushima, Haruko Kubo

**Kumamoto University, Japan**

Terumasa Ikeda, Chihiro Motozono, Hesham Nasser, Ryo Shimizu, Yue Yuan,

Kazuko Kitazato, Haruyo Hasebe, Takamasa Ueno

**University of Miyazaki, Japan**

Akatsuki Saito, Erika P Butlertanaka, Yuri L Tanaka

**National Institute of Infectious Diseases, Japan**

Kenzo Tokunaga, Seiya Ozono

**Tokyo Metropolitan Institute of Public Health, Japan**

Kenji Sadamasu, Hiroyuki Asakura, Isao Yoshida, Mami Nagashima, Kazuhisa Yoshimura

**Ecuador-COVID19 Consortium**

**Universidad San Francisco de Quito, COCIBA, Instituto de Microbiología, Ecuador**

Sully Márquez, Belén Prado-Vivar, Mónica Becerra-Wong, Mateo Caravajal, Gabriel Trueba, Patricio Rojas- Silva

**Universidad San Francisco de Quito, COCSA, Escuela de Medicina, Ecuador**

Michelle Grunauer

**Universidad San Francisco de Quito, COCIBA, Laboratorio de Biotecnología Vegetal, Ecuador**

Bernardo Gutierrez, Juan José Guadalupe

**Laboratorio INTERLAB, Ecuador**

Juan Carlos Fernández-Cadena

**Universidad Espíritu Santo, Laboratorio de Omicas, Ecuador**

Derly Andrade-Molina

**Universidad Internacional del Ecuador, Facultad de Ciencias Médicas, de la Salud y la Vida, Ecuador**

Manuel Baldeon

**Centros Médicos Dr. Marco Albuja, Ecuador**

Andrea Pinos

**Acknowledgments**

We would like to thank all members of The Genotype to Phenotype Japan (G2P-Japan) and Ecuador-COVID19 Consortia. We thank Dr. Kenzo Tokunaga (National Institute of Infectious Diseases, Japan) for sharing materials.

This study was supported in part by AMED Research Program on Emerging and Re-emerging Infectious Diseases 20fk0108146 (to Kei Sato), 20fk0108270 (to Kei Sato), 20fk0108413 (to Kei Sato) and 20fk0108451 (to G2P-Japan Consortium, Akifumi Takaori-Kondo, and Kei Sato); AMED Research Program on HIV/AIDS 21fk0410039 (to Kotaro Shirakawa and Kei Sato); JST SICORP (e-ASIA) JPMJSC20U1 (to Kei Sato); JST SICORP JPMJSC21U5 (to Kei Sato), JST CREST JPMJCR20H4 (to Kei Sato); JSPS KAKENHI Grants-in-Aid for Scientific Research B 18H02662 (to Kei Sato) and 21H02737 (to Kei Sato); JSPS Fund for the Promotion of Joint International Research (Fostering Joint International Research) 18KK0447 (to Kei Sato); JSPS Core-to-Core Program JPJSCCA20190008 (A. Advanced Research Networks) (to Kei Sato); The Tokyo Biochemical Research Foundation (to Kei Sato); and Joint Usage/Research Center program of Institute for Frontier Life and Medical Sciences, Kyoto University (to Kei Sato).
